# hei-tag: a highly efficient tag to boost targeted genome editing

**DOI:** 10.1101/2021.05.27.445956

**Authors:** Thomas Thumberger, Tinatini Tavhelidse, Jose Arturo Gutierrez-Triana, Rebekka Medert, Alex Cornean, Bettina Welz, Marc Freichel, Joachim Wittbrodt

**Affiliations:** Centre for Organismal Studies (COS), Heidelberg University, Im Neuenheimer Feld 230, D-69120 Heidelberg, Germany; Heidelberg Biosciences International Graduate School (HBIGS), Heidelberg, Germany; Escuela de Microbiología, Facultad de Salud, Universidad Industrial, Santander, Colombia; Institute of Pharmacology, Heidelberg University, Heidelberg, Germany; DZHK (German Centre for Cardiovascular Research), partner site Heidelberg/Mannheim, Heidelberg, Germany

## Abstract

Precise, targeted genome editing by CRISPR/Cas9 is key for basic research and translational approaches in model and non-model systems^1^. While active in all species tested so far, editing efficiencies still leave room for improvement. To reach its target, the bacterial Cas9 needs to be efficiently shuttled into the nucleus as attempted by fusion of nuclear localization signals (NLSs) to the Cas9 protein^2^. Additional domains such as FLAG- or myc-tags are added for immediate detection or straight-forward purification^3^. To avoid steric hinderance impacting on activity, amino acid linkers are employed connecting Cas9 and additional domains.

We present the ‘hei-tag (<u>h</u>igh <u>e</u>ff<u>i</u>ciency-tag)’, boosting the activity of the wide variety of CRISPR/Cas genome editing tools. The addition of the hei-tag to Cas9 or a C-to-T base editor dramatically enhances the respective targeting efficiency in model systems ranging from fish to mammals, including tissue culture applications. This allows to instantly upgrade existing and potentially highly adapted systems as well as establish novel highly efficient tools.

## Results and Discussion

For nuclear localization of proteins of interest like the Cas9 enzyme, the monopartite NLS originating from the SV40 large T-antigen^4^ or a bipartite NLS discovered in *Xenopus* nucleoplasmin are routinely employed^5^.

For high targeting efficiency with low mosaicism, a peak activity should be achieved in the zygote or at early cleavage stages. However, the nuclear localization activity of commonly used NLSs is tightly controlled during early development^6^ and is first detectable during gastrulation. In fish embryos, an optimized artificial NLS^7^ (oNLS) facilitates prominent nuclear localization already immediately after fertilization, while the SV40 NLS acts most prominently much later and facilitates nuclear localization approximately at the 1000 cell stage.

Assessing the efficiency of targeted genome editing requires a reliable and quantitative readout based on an apparent phenotype. We established a quantitative assay for loss-of-eye-pigmentation to address the activity of different Cas9 variants in two teleost model systems, medaka (*Oryzias latipes*) and zebrafish (*Danio rerio*) covering a wide evolutionary distance of 200 million years^8^. Our assay on retinal pigmentation provides a highly reproducible quantitative readout for the loss of the conserved transporter protein *oculocutaneous albinism type 2 (oca2),* required for melanin biosynthesis (Fig. 1a). Only its bi-allelic inactivation results in the loss of pigmentation of eyes and skin^9^. Altered pigmentation in the eye thus provides a quantitative readout for bi-allelic targeting efficiency, while the degree of mosaicism reflects the time point of action (uniform-early, mosaic-late).

**Fig. 1:**
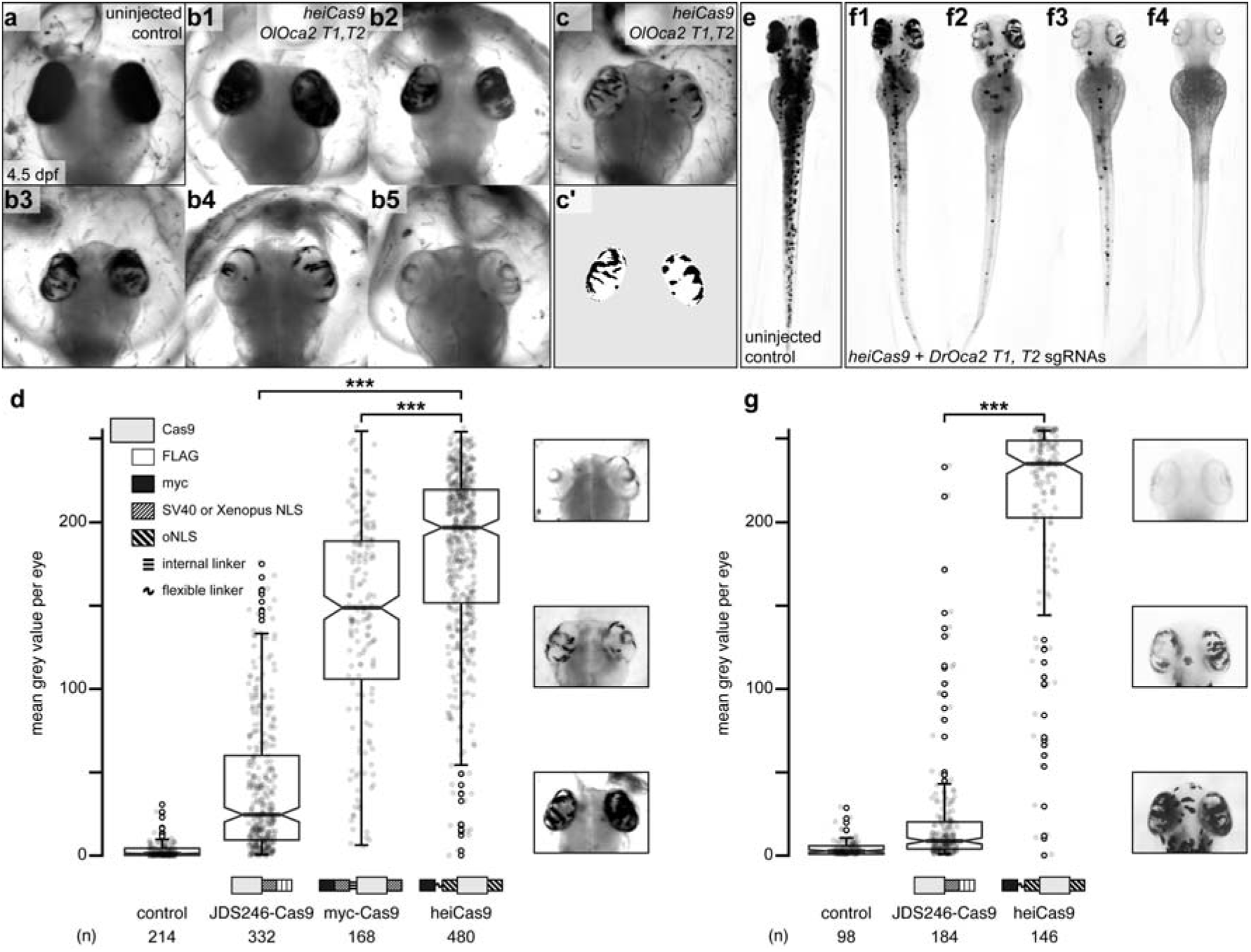
heiCas9 exhibits outstanding bi-allelic targeting activity in fish. Phenotypic range and quantification of *OlOca2 T1, T2* and *DrOca2 T1, T2* sgRNAs/*Cas9 variant* mediated loss of pigmentation in medaka (a-d) and zebrafish (e-g). (a) Fully pigmented eyes in uninjected control medaka embryo at 4.5 dpf. (b1-b5) Range of typically observed loss-of-pigmentation phenotypes upon injection with *heiCas9* mRNA and *OlOca2 T1, T2* sgRNAs. The observed phenotypes range from almost full pigmentation (b1) to completely unpigmented eyes (b5). (c) Minimum intensity projection of a medaka embryo at 4.5 days after injection with *heiCas9* and *OlOca2 T1, T2* sgRNAs. (c’) Locally thresholded pigmentation on elliptical selection per eye (same embryo as in c). (d) Quantification of mean grey values (0 = fully pigmented, 255 = completely unpigmented) of individual eyes from Oca2 knock-out medaka embryos co-injected with *OlOca2 T1*, *T2* sgRNAs and mRNAs of *JDS246-Cas9, myc-Cas9* and *heiCas9* respectively. Medians: uninjected control = 0.75; JDS246-Cas9 = 24.60; myc-Cas9 = 148.81; heiCas9 = 196.94. Note: highly significant pigment loss (8-fold increase) in heiCas9 versus JDS246-Cas9 crispants (p = 7.7e-113) and versus myc-Cas9 (p = 2.3e-13). (e) Fully pigmented uninjected control zebrafish embryo at 2.5 dpf. (f1-f4) Range of typically observed loss-of-pigmentation phenotypes upon injection with *heiCas9* mRNA and *DrOca2 T1, T2* sgRNAs. The observed phenotypes range from almost full pigmentation (f1) to completely unpigmented eyes and body (f4). (g) Quantification of mean grey values of individual eyes from Oca2 knock-out zebrafish embryos co-injected with *DrOca2 T1, T2* sgRNAs and mRNAs of *JDS246-Cas9* and *heiCas9* respectively. Medians: uninjected control = 2.53; JDS246-Cas9 = 8.60; heiCas9 = 234.54. Note the highly significant pigment loss (27-fold increase) in heiCas9 versus JDS246-Cas9 crispants (p = 5.3e-48). dpf, days post fertilization; mean grey values ranged from 0, i.e. fully pigmented eye to 255, i.e. complete loss of pigmentation; n, number of eyes analyzed. Statistical analysis performed in R, pairwise Wilcoxon rank sum test, Bonferroni corrected.

To facilitate uniform Cas9 action, we followed our successful mRNA injection protocol^10^. One-cell stage medaka embryos were co-injected with sgRNAs targeting the *oca2* gene (*OlOca2 T1, T2*) together with mRNA encoding the respective Cas9 variant. Injected embryos were fixed well after the onset of pigmentation at 4.5 days post fertilization^11^ and subjected to image analysis (Fig. 1b). In brief, the eyes were segmented, (residual) pigmentation was thresholded (Fig. 1c-c’) and quantified according to mean grey values (0, i.e. fully pigmented, 255, i.e. completely unpigmented, Fig. 1d).

We first established the base activity level for the assay and determined the activity of one of the earliest Cas9 variants publicly available, a Cas9 carrying a C-terminal SV40 NLS followed by three FLAG tags (JDS246-Cas9, Plasmid #43861 Addgene). The analysis of medaka embryos injected with *JDS246-Cas9* revealed few bi-allelic inactivation events of the *oca2* gene as apparent by patchy unpigmented domains in the eye (median of mean grey values = 24.60 compared to uninjected controls, median = 0.75; Fig. 1d). This mosaic distribution of small, unpigmented areas indicated that targeting occurred only in few cells at blastula or later stages of development.

We next analysed the activity of an improved myc-Cas9 variant in which a SV40 NLS and a *Xenopus* nucleoplasmin bipartite NLS are flanking the Cas9 enzyme at the N- and C-terminus, respectively^3^. The N-terminal SV40 NLS is preceded by a myc-tag and connects to Cas9 via a VPAA linker, an amino acid spacer regarded as rigid^12^. In comparison to JDS246-Cas9, this variant showed a 6-fold enhanced activity as reflected by the increase in mean grey values (median = 148.81; Fig. 1d). In addition, targeting in earlier embryonic stages (before and around blastula stages) was apparent by larger clones and a lower degree of mosaicism.

Different tags fused to the same enzyme apparently impact differentially on its resulting activity. We combined the early prominent nuclear localization mediated by the oNLS^7^ and reduced steric hinderance of the fused domains^12^ by flexible linkers to further increase gene editing efficiency of Cas9.

We therefore developed the hei-tag for enhancing the activity of proteins acting in the nucleus. The hei-tag is composed of a myc-tag connected via a flexible linker to an oNLS at the N-terminus complemented by a second oNLS fused to the C-terminus of the tagged protein, initially a mammalian codon optimized *Cas9* variant (see Supplementary Data for sequence).

When assessing the activity of the resulting heiCas9 we detected an 8-fold increase in bi-allelic targeting efficiency compared to JDS246-Cas9 (median = 196.94; Fig. 1d). Embryos co-injected with this *“heiCas9”* mRNA and sgRNAs against *oca2* showed almost no residual pigmentation, and in comparison to myc-Cas9 the activity of heiCas9 was increased to 132%. Using heiCas9, the rate of mosaicism was by far the lowest, arguing for an early time point of action due to high activity and efficient nuclear translocation of the tagged heiCas9 variant already at the earliest cleavage stages.

To address whether the enhancement by hei-tag fusion to Cas9 variants is widely applicable to different models, we next compared the activities of the JDS246-Cas9 and heiCas9 in a second, evolutionarily distant fish species *Danio rerio* (zebrafish) targeting the orthologous *oca2* gene (sgRNAs *DrOca2 T1, T2*). Injected and control embryos were fixed well after the onset of pigmentation at 2.5 dpf^13^ (Fig. 1e-f) and subjected to the quantitative assay for eye pigmentation described above. Taking the activity of JDS246-Cas9 as basal level (median = 8.60), heiCas9 delivered an outstanding targeting efficiency (median = 234.54), reflecting a significant (p<0.001), 27-fold increase (Fig. 1g). Similar to the results in medaka, also here highly efficient, early targeting was achieved as reflected by a massive reduction of the degree of mosaicism.

Taken together, addition of the hei-tag to a mammalian codon optimized Cas9 resulted in the highly efficient heiCas9, which allows for an up to 27-fold increase in targeting efficiency. It prominently inactivates both alleles of the targeted *oca2* locus, with an early onset of action upon injection of *heiCas9* mRNA and the respective sgRNAs at the one-cell stage.

While the early onset of action is required for uniform editing in developing organisms, cell culture approaches demand efficient translocation of the sgRNA/Cas9 complex in a large number of cells. To validate the range of action on the one hand and to address the relevance of the hei-tag in a mammalian setting, we expanded the scope of the analysis to mammalian cell culture. We focused on mRNA-based assays and compared the activity of heiCas9 to state of the art Cas9 variants, i.e. the mRNAs of *GeneArt^®^ CRISPR nuclease* as well as *JDS246-Cas9* in mouse SW10 cells. We assessed the respective genome editing efficiencies by independent and complementary tools, the Tracking of Indels by Decomposition (TIDE) analysis^14^ as well as by Inference of CRISPR Editing (ICE)^15^. Both approaches decompose the mixed Sanger reads of PCR products spanning the CRISPR target site and compute an efficiency score as well as the distribution of expected indels. Mouse SW10 cells were co-transfected with *MmPrx* crRNA / ATTO-550-linked tracrRNA and the mRNAs of either *GeneArt^®^ CRISPR nuclease, JDS246-Cas9* or *heiCas9.* The targeted *Periaxin (Prx*) locus was PCR amplified and sequenced. Similar to targeting *in organismo*, heiCas9 also exhibited the highest genome editing efficiency when compared to GeneArt^®^ CRISPR nuclease (TIDE: 123.1%, ICE: 111%) and JDS246-Cas9 (TIDE: 123.6%, ICE: 113%) in mammalian cell culture (Fig. 2, Supplementary Fig. 1, R^2^ >0.9 (TIDE) and >0.9 (ICE) for all mRNAs tested). Notably, the KO-score efficiencies (ICE) amounted to 167% compared to GeneArt^®^ CRISPR nuclease and to 173% compared to JDS246-Cas9, indicating higher abundance of frameshifts^15^.

**Fig. 2:**
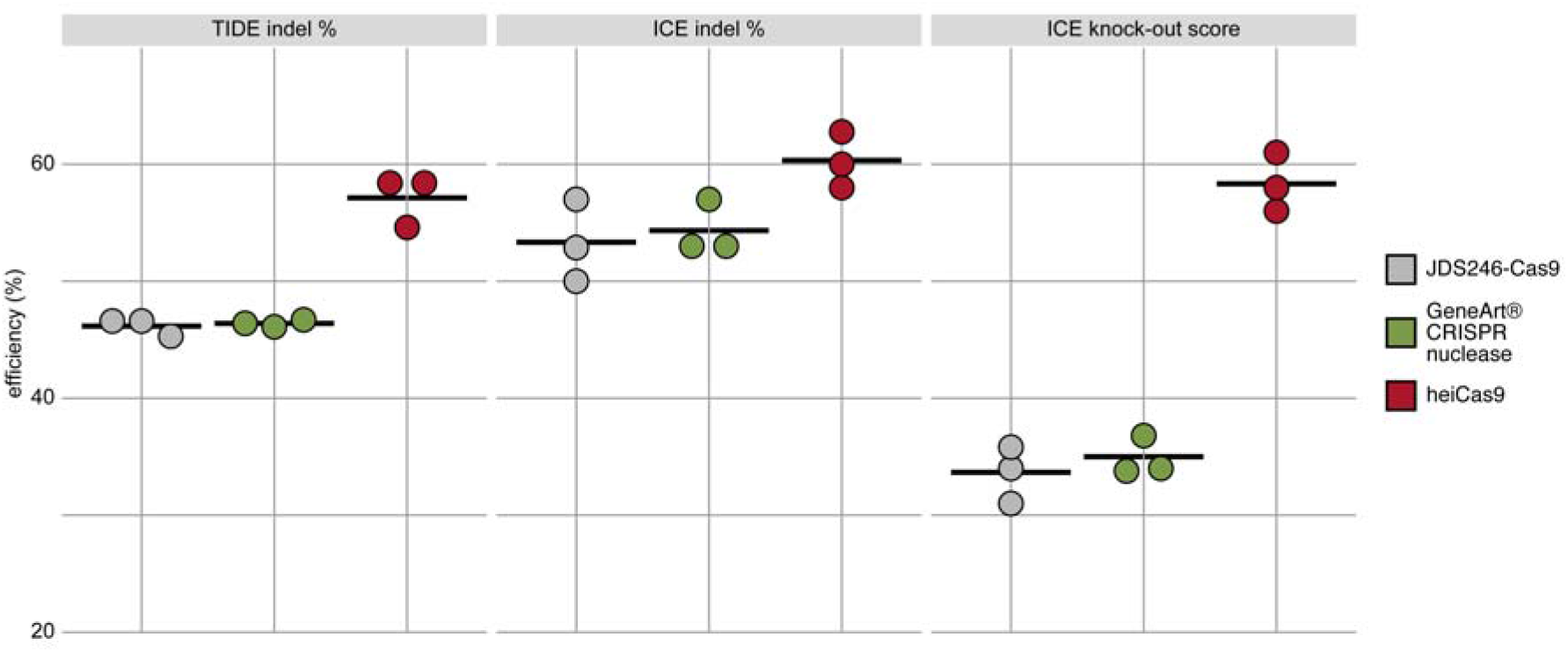
heiCas9 consistently exhibits high genome editing efficiency in mammalian cells. Mouse SW10 cells were co-transfected with *MmPrx* crRNA and mRNAs of *JDS246-Cas9, GeneArt^®^ CRISPR nuclease* and *heiCas9* respectively. Genome editing efficiency was assessed by TIDE and ICE tools. ICE knock-out score represents proportion of indels that indicate a frameshift or ≥ 21 bp deletion. Data points represent three biological replicates, black line indicates respective mean: TIDE indel %: JDS246-Cas9 = 46.2; GeneArt^®^ CRISPR nuclease = 46.4, heiCas9 = 57.1; ICE indel %: JDS246-Cas9 = 53.3; GeneArt^®^ CRISPR nuclease = 54.3, heiCas9 = 60.3; ICE knock-out score %: JDS246-Cas9 = 33.7; GeneArt^®^ CRISPR nuclease = 35.0, heiCas9 = 58.3. R^2^ >0.9 (TIDE) and >0.9 (ICE) for all mRNAs tested. For representative indel spectrum and quality control diagrams for each mRNA see Supplementary Fig. 1.

Remarkably, *heiCas9* transfected cells showed a highly increased number of mutant alleles with an increased abundance of a 26 nt deletion when compared to GeneArt^®^ CRISPR nuclease and JDS246-Cas9 (Supplementary Fig. 1).

Given the observed boosting of Cas9 activity by the addition of the hei-tag, we next tested if addition of the hei-tag also improves further Cas9-based techniques. We fused the hei-tag to the cytosine (C-to-T) base editor BE4-Gam^16^. In base editors (BEs), a modified Cas9, that does not introduce a double-strand break (Cas9 nickase or Cas9n) is employed together with a nucleobase deaminase for precisely targeted nucleotide editing^17^. Here, we tested the hei-tag in combination with BE4-Gam^16^ to introduce a pre-mature STOP codon in exon 9 of the *oca2* open reading frame (ORF) by transition of cytosine 997 to thymine (C997T, leading to Q333*). Again, the loss of pigmentation was used as proxy for bi-allelic targeting efficiency. Medaka one-cell stage embryos were injected with one of three sgRNAs (*OlOca2 T1, T3* or *T4*) as well as with either *BE4-Gam* or *heiBE4-Gam* and screened for pigmentation phenotypes at 4.5 dpf. For each sgRNA, loss of pigmentation was much more pronounced when combined with heiBE4-Gam rather than BE4-Gam (Fig. 3a). Quantification of Sanger sequencing reads confirmed an increase of all C-to-T transitions (44.2% ± 6.8% for BE4-Gam vs. 74.1% ± 8.9% for heiBE4-Gam; Supplementary Fig. 2). In particular the C997T transition introducing a pre-mature STOP codon was increased 1.7-fold (i.e. 41% in BE4-Gam to 68% in heiBE4-Gam) in case of *heiBE4-Gam* (Fig. 3b, c). Notably, in *heiBE4-Gam* injections, for each of the three cytosines in the base editing window, the C-to-T transition rate was higher than 60%, a level never observed in *BE4-Gam* injected embryos (Supplementary Fig. 2).

**Fig. 3:**
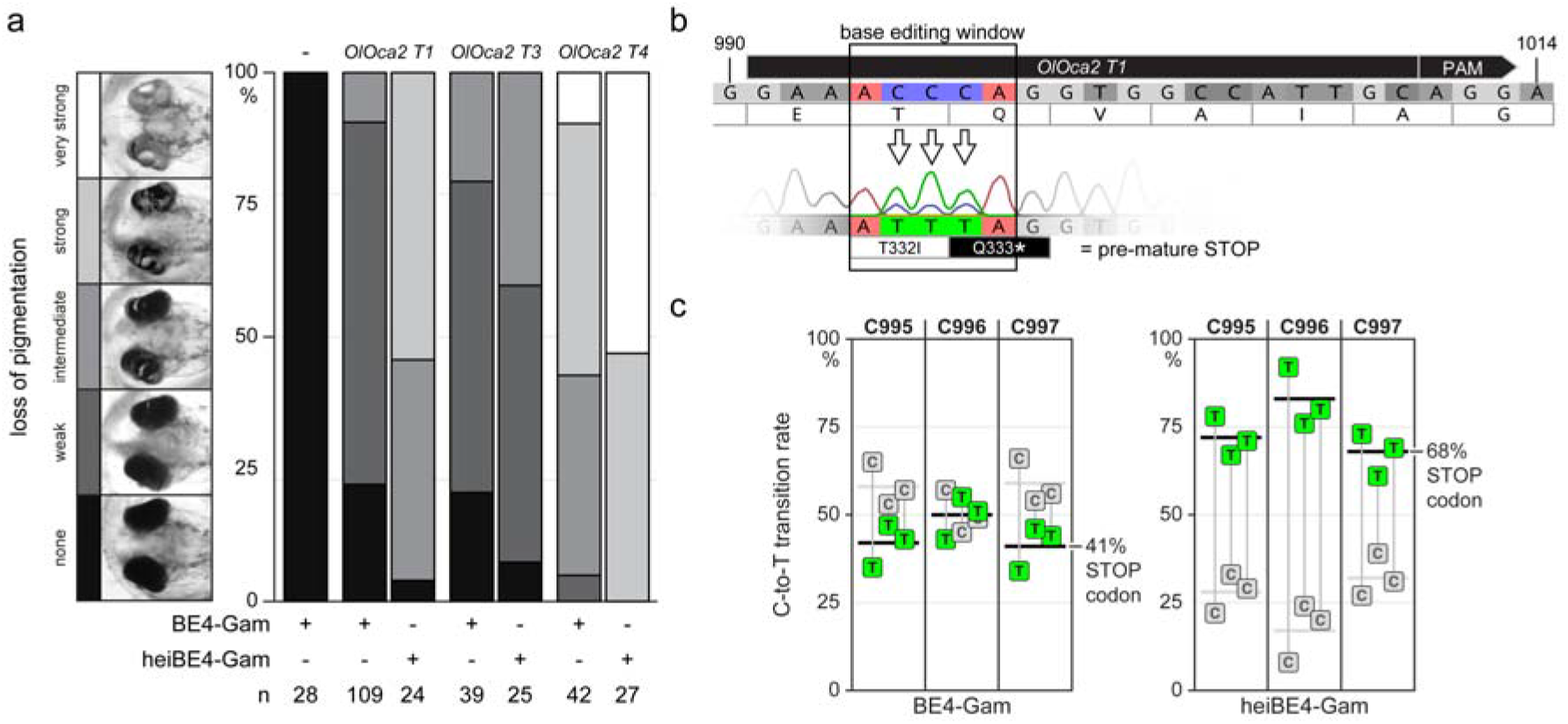
heiBE4-Gam mediates highly efficient cytosine to thymine transition in medaka embryos. Phenotypic range and quantification of heiBE4-Gam mediated cytosine to thymine transitions in medaka embryos. (a) Categories of typically observed loss-of-pigmentation phenotypes upon injection with *BE4-Gam* or *heiBE4-Gam* and *OlOca2 T1, T3, T4* sgRNAs. The observed pigmentation phenotypes range from (almost) unpigmented eyes, i.e. a very strong knock-out (top panel; white) over intermediate (central panel; grey) to no loss-of-pigmentation (bottom panel; black). Quantification of phenotype categories resulting from injections with either *BE4-Gam* or *heiBE4-Gam* and *OlOca2 T1, T3,* or *T4* sgRNAs. Note: dramatic increase of bi-allelic knock-out rate when using *heiBE4-Gam.* n, number of embryos analyzed. (b) Schematic representation of base editing window in *OlOca2 T1* target site. C-to-T transition of C995 and C996 edits the threonine (T) codon to isoleucine (I) (T332I); C997T creates a pre-mature STOP codon (Q333*). Nucleotide positions refer to open reading frame. (c) Quantification of Sanger sequencing reads at nucleotides C995, C996, C997 inside the base editing window of three injected embryo pools reveals overall dramatic increase of C-to-T base transition when using heiBE4-Gam. Note 1.7-fold increase of C997T transition, i.e. efficient introduction of a pre-mature STOP codon. Mean values indicated by thick horizontal lines, cf. Supplementary Figure 2.

In conclusion, using the hei-tag to extend the ORFs of a mammalian codon optimized SpCas9 or a C-to-T base editor (BE4-Gam) we dramatically improved genome targeting efficiency.

The resulting heiCas9 represents a highly efficient endonuclease overcoming the limitations of current *SpCas9* variants by dramatically increasing the efficiency of bi-allelic targeting (up to 27-fold) in two evolutionarily distant fish models. Notably, heiCas9 markedly increased the bi-allelic targeting rate in both, medaka and zebrafish, indicating a high targeting efficiency already at the earliest stages of development.

Strikingly, the hei-tag not only showed unprecedented efficiency *in organismo*, but also delivered highest targeting efficiencies in mammalian cell culture (1.7-fold increase in mouse SW10 cells) when fused to Cas9 and compared to commercially available *SpCas9* mRNAs.

In line with those findings, the simple addition of the hei-tag sequence also potentiated the activity of a cytosine base editor, with heiBE4-Gam resulting in an overall increase of about 30% of C-to-T transition rates (Fig. 3 and Supplementary Fig. 2).

Taken together, the boosting activity of the hei-tag is neither limited by the species nor the approach, making it a powerful tweak to swiftly upgrade any specifically adapted Cas-based genome editing approach^17^.

## Materials and Methods

### Fish maintenance

Zebrafish (*Danio rerio*) and medaka (*Oryzias latipes*) fish were bred and maintained as previously described^18,19^. The animal strains used in the present study were zebrafish AB/back and medaka Cab. All experimental procedures were performed according to the guidelines of the German animal welfare law and approved by the local government (Tierschutzgesetz §11, Abs. 1, Nr. 1, husbandry permit number 35–9185.64/BH Wittbrodt).

### Plasmids

The mammalian codon-optimized (Geneious 8.1.9 (https://www.geneious.com)) *Cas9* sequence was gene-synthesized (GeneArt, ThermoFisher Scientific) as template for cloning *heiCas9* using primers (Table 1) containing the sequences coding for the hei-tag (myc-tag (EQKLISEEDL), flexible linker (GGSG) and an optimized oNLS (PPPKRPRLD)^7^ (Supplementary Data). Cloning into the pCS2+ plasmid^20^ (multiple cloning site extended for *AgeI* site downstream of *BamHI* site) was performed using *AgeI* and *XbaI* restriction sites included in the 5’ region of the forward or reverse primers, respectively. See Supplementary Data for full sequence of *heiCas9.* For consistent mRNA synthesis, the published *myc-Cas9^3^* was re-established with the pX330-U6-Chimeric_BB-CBh-hSpCas9 vector as template, primer based exchange of the N-terminal FLAG tag with the myc-tag sequence and brought into pCS2+^20^ using *AgeI* and *XbaI* restriction sites included in the 5’ region of the respective primers as well. pX330-U6-Chimeric_BB-CBh-hSpCas9 was a gift from Feng Zhang (Addgene plasmid #42230)^2^.

**Table 1.**
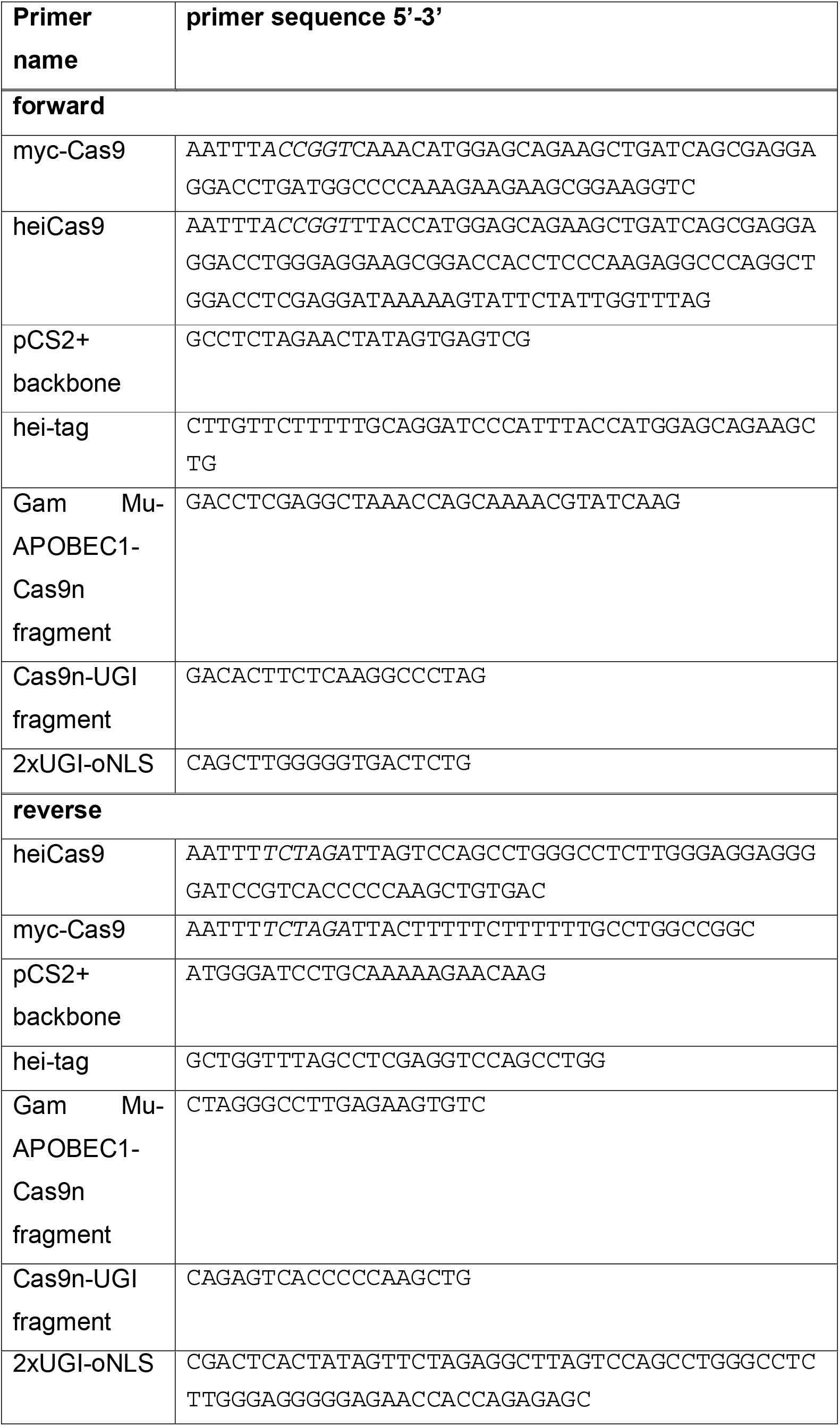
Primer sequences used in this study. Restriction enzyme sites used for cloning into pCS2+ plasmid are indicated in italics (*AgeI* in the forward primer, *XbaI* in the reverse primer).

BE4-Gam was subcloned from pCMV(BE4-Gam) (Addgene plasmid #100806, a gift from David Liu)^16^ in a two step-process, first into pJET1.2 (Thermo Scientific), then into pGGEV4^21^ (Addgene plasmid #49284), by *BamHI, EcoRV* and *KpnI* restriction sites to create pGGEV4(BE4-Gam). heiBE4-Gam was assembled into pCS2+^20^ by NEBuilder^®^ HiFi DNA Assembly (NEB) with four inserts using Q5 polymerase PCR products (NEB): pCS2+ backbone, hei-tag, Gam Mu-APOBEC1-Cas9n fragment, Cas9n-UGI fragment, 2xUGI-oNLS (see Table 1 for primers used).

*Oca2 sgRNAs* for medaka (*OlOca2*) and zebrafish (*DrOca2*) were designed using the CCTop target predictor with standard parameters^22^. The following target sites were used [PAM in brackets]: *OlOca2 T1* (GAAACCCAGGTGGCCATTGC[AGG]), *OlOca2 T2* (TTGCAGGAATCATTCTGTGT[GGG]), *OlOca2 T3* (GATCCAAGTGGAGCAGACTG[AGG]), *OlOca2 T4* (CACAATCCAGGCCTTCCTGC[AGG]) *DrOca2 T1* (GTACAGCGACTGGTTAGTCA[TGG]) and *DrOca2 T2* (TAAGCACGTAGACTCCTGCC[AGG]). *OlOca2 T1, OlOca2 T2* and *DrOca2 T1* were the same as in Hammouda et al., 2019^23^, *OlOca2 T3* was the same as in Lischik et al., 2019^9^ (OCA2_4). Cloning of sgRNA templates was performed as described^22^. Plasmid DR274 was a gift from Keith Joung (Addgene plasmid #42250).

### *In vitro* transcription of mRNA

All pCS2+ constructs in this work were linearized using NotI-HF (NEB), pGGEV4(BE4-Gam) was linearized using SpeI-HF (NEB) and mRNA was transcribed *in vitro* using the mMESSAGE mMACHINE SP6 transcription kit (ThermoFisher Scientific, AM1340). The *JDS246-Cas9* was linearized with MssI FD (ThermoFisher Scientific) and transcribed *in vitro* using the mMESSAGE mMACHINE T7 Ultra Transcription Kit (ThermoFisher Scientific, AM1345). JDS246-Cas9 was a gift from Keith Joung (Addgene plasmid #43861). *Oca2* sgRNAs were synthesized using the MEGAscript T7 transcription kit (Thermo Fisher Scientific, AM1334) after plasmid digestion with DraI FD (ThermoFisher Scientific).

### Microinjection

Zebrafish and medaka zygotes were injected with *Cas9* mRNA at either 30 ng/μl or 150 ng/μl, *Oca2* sgRNAs at 30 ng/μl and *H2B-GFP* mRNA at 10 ng/μl as an injection marker. For the base editing experiments, medaka zygotes were injected with *BE4-Gam* or *heiBE4-Gam* mRNA at 50 or 150 ng/μl, *Oca2* sgRNA at 30 ng/μl and *GFP* mRNA at 20 ng/μl as injection tracer. Injected embryos were maintained at 28°C in zebrafish medium^19^ or medaka embryo rearing medium (ERM, 17 mM NaCl; 40 mM KCl; 0.27 mM CaCl_2_•2H_2_O; 0.66 mM MgSO_4_•7H_2_O,17 mM Hepes).

Embryos were screened for *GFP* expression seven hours or one day after injection, and GFP negative specimens were discarded.

### Image acquisition and phenotype analysis

Medaka 4.5 days post fertilization (dpf) embryos^11^ and zebrafish 2.5 dpf^13^ embryos were fixed with 4% paraformaldehyde in 2xPTW (10x PBS pH 7.3, 20% Tween20). Images of medaka embryos were acquired with the high content screening ACQUIFER Imaging Machine (DITABIS AG, Pforzheim, Germany). Images of zebrafish embryos were acquired with a Nikon digital sight DS-Ri1 camera mounted onto a Nikon Microscope SMZ18 and the Nikon Software NIS-Elements F version 4.0. Only properly developed embryos were included in the following analysis. Image analysis was performed with Fiji^24^, i.e. mean grey values were obtained on minimum intensity projections and locally thresholded (Phansalkar algorithm with parameters r = 20, p = 0.4, k = 0.4) pictures and elliptical selections for each individual eye. The mean grey value per eye was used for the boxplot and statistical analysis (pairwise comparisons using Wilcoxon rank sum test, Bonferroni corrected) in RStudio^25^. For the base editing experiments, pigmentation phenotypes were scored 4.5 days after injection on properly developed embryos.

### Genotyping

Pools of 5 randomly selected embryos were used to prepare DNA for genotypic validation of base editing by lysis in DNA extraction buffer (0.4 M Tris/HCl pH 8.0, 0.15 M NaCl, 0.1% SDS, 5 mM EDTA pH 8.0, 1 mg/ml proteinase K) at 60°C overnight. Proteinase K was inactivated at 95°C for 10 min and the solution was diluted 1:2 with H_2_O.

Genotyping was performed with Q5 polymerase (NEB), primers fwd 5’-GTTAAAACAGTTTCTTAAAAAGAACAGGA-3’ and rev 5’-AGCAGAAGAAATGACTCAACATTTTG-3’ (annealing at 62°C) on 1 μl of diluted DNA sample according to the manufacturer’s instructions with 30x PCR cycles. PCR products were analyzed on a 1% agarose gel, bands excised, DNA-extraction performed using innuPREP Gel Extraction Kit (Analytik Jena) according to the manufacturer’s instructions and subjected to sequencing.

### Cell lines

Mouse SW10 cells (ATCC, CRL-2766) were cultured in DMEM (Gibco) supplemented with 1 g/ml glucose containing 10% FCS (Sigma), 1% penicillin (10,000 units/ml; Gibco) and 1% streptomycin (10 mg/ml; Gibco) and maintained at 33°C and 5% CO_2_. Cells were passaged at 80-90% confluency. 24 h before transfection cells were seeded in a density of 85,000 cells per 12-well.

### CRISPR Transfection

crRNA targeting exon 6 (TCGTATCCAGACACCGTCCC[GGG], PAM in brackets) of the mouse *Periaxin (MmPrx*) gene was selected from the IDT (crRNA XT) predesign crRNA database. crRNA (50 μM) and Alt-R^®^ CRISPR-Cas9 tracrRNA, ATTO-550 (50 μM; IDT, 1075927) were diluted in nuclease-free duplex buffer (IDT) to a final concentration of 1 μM and incubated at 95°C for 5 minutes. 1 μg of the corresponding *Cas9* mRNA (*GeneArt^®^ CRISPR nuclease* Invitrogen, A29378; *JDS246-Cas9* or *heiCas9*) and 15 μl of tracrRNA+crRNA Mix (1 μM) were diluted in 34 μl Opti-MEM I (Gibco) and mixed with 3 μl Lipofectamine RNAiMAX (ThermoFisher) diluted in 47 μl Opti-MEM I. The tracrRNA+crRNA lipofection mix was incubated for 20 min at RT. Cell culture medium was exchanged with 900 μl Opti-MEM I and the tracrRNA+crRNA lipofection mix was added dropwise to the well. After 48h, genomic DNA was extracted using the DNeasy Blood and Tissue Kit (Qiagen, 69506) following the manufacturer’s protocol. Q5-PCR was carried out using primers flanking the targeted exon 6 (fwd 5’-GAGACACTCACTCCAGACCC-3’; rev 5’-ACTCAGTAACCCAACAGCCA-3’) and 30 cycles. PCR amplicons were purified using the Monarch DNA Gel Extraction Kit (NEB, T1020S) and subjected to sequencing.

### Sequencing

Sanger sequencing was performed by Eurofins Genomics using fwd 5’-GTTAAAACAGTTTCTTAAAAAGAACAGGA-3’ to evaluate base editing and using fwd 5’-GAGACACTCACTCCAGACCC-3’ and rev 5’-ACTCAGTAACCCAACAGCCA-3’ to evaluate genome editing in SW10 cells. Quantification of base editing from sanger sequencing reads was performed with EditR^26^. Genome editing efficiency was assessed by sequence analysis using the TIDE web tool^14^ and by ICE^15^ using default parameters.

### Data visualisation

Data visualisation and figure assembly was performed using Fiji^24^, ggplot2^27^ in RStudio^25^, Geneious Prime 2019.2.1 and Adobe^®^ Illustrator^®^ CS6.

## Acknowledgements

This research was funded through the German Science funding agency (DFG, CRC873, project A3 and FOR2509 project 10 (WI 1824/9-1) to J.W. and CRC1118, project S03 to M.F., European Research Council (GA 294354-ManISteC to J.W.)). A.C., B.W., R.M. and T.Ta. are members/alumni of HBIGS, the Heidelberg Biosciences International Graduate School. B.W. was supported by the Deutsches Zentrum für Herz-Kreislauf-Forschung (DZHK B20-024 SE). We thank T. Kellner for sgRNA and Cas9 mRNA synthesis. We are thankful to M. Majewski, E. Leist and A. Saraceno for fish husbandry. We thank all members of the Wittbrodt lab for their critical, constructive feedback on the procedure and the manuscript.

## Author contributions

T.T., T.Ta., J.A.G.T. and J.W. conceived the study and designed the experiments; J.A.G.T. cloned the *Cas9* variants, T.T. performed the *Cas9* injection experiments in fish embryos, R.M. designed and performed cell culture experiments, A.C. designed and analyzed the base editing experiments he performed together with B.W., T.T. and T.Ta. quantified results in fish embryos; T.Ta., T.T. and J.W. wrote the manuscript with contributions from R.M., A.C., B.W. and M.F.; M.F. and J.W. supervised and J.W. led the project.

