## Supplementary Data and Figures for "hei-tag: a highly efficient tag to boost targeted genome editing"

Nucleotide and translated amino acid sequence of heiCas9.

**myc**-flexible linker-**oNLS**-Cas9-**oNLS**

1 atggagcagaagctgatcagcgaggaggacctgggaggaagcggaccacctcccaagagg 60
1 M **E Q K L I S E E D L** G G S G **P P P K R** 20
61 cccaggctggacctcgaggataaaaagtattctattggtttagacatcggcaccaacagc 120
21 **P R L D** L E D K K Y S I G L D I G T N S 40
121 gtgggctgggccgtgatcaccgacgagtacaaggtgcccagcaagaaattcaaggtgctg 180
41 V G W A V I T D E Y K V P S K K F K V L 60
181 ggcaacaccgacagacacagcatcaagaaaaacctgatcggcgccctgctcttcgactcc 240
61 G N T D R H S I K K N L I G A L L F D S 80
241 ggcgaaaccgccgaggccaccagactgaagagaaccgccagacggagatacaccagacgg 300
81 G E T A E A T R L K R T A R R R Y T R R 100
301 aagaatagaatctgctacctgcaggagatcttcagcaacgagatggccaaggtggacgat 360
101 K N R I C Y L Q E I F S N E M A K V D D 120
361 agcttctttcacagactggaagagagcttcctggtggaagaggacaagaaacacgagaga 420
121 S F F H R L E E S F L V E E D K K H E R 140
421 caccccatcttcggcaacatcgtggacgaggtggcctaccacgagaagtaccccaccatc 480
141 H P I F G N I V D E V A Y H E K Y P T I 160
481 taccacctgagaaagaaactggtggacagcaccgacaaggccgacctgagactgatctac 540
161 Y H L R K K L V D S T D K A D L R L I Y 180
541 ctggcactggcccacatgatcaagttcagaggccacttcctgatcgagggcgacctgaac 600
181 L A L A H M I K F R G H F L I E G D L N 200
601 cccgacaacagcgacgtggacaagctgttcatccagctggtgcagacctacaaccagctg 660
201 P D N S D V D K L F I Q L V Q T Y N Q L 220
661 ttcgaagagaaccctatcaacgccagcggcgtggacgccaaggccatcctgagcgccaga 720
221 F E E N P I N A S G V D A K A I L S A R 240
721 ctcagcaagagcagacggctggagaacctgatcgcccagctgcccggcgagaagaaaaac 780
241 L S K S R R L E N L I A Q L P G E K K N 260
781 ggcctgttcggcaacctgatcgccctgagcctgggcctgacccccaacttcaagagcaac 840
261 G L F G N L I A L S L G L T P N F K S N 280
841 ttcgacctggccgaggacgccaagctgcagctgagcaaggacacctacgacgatgacctg 900
281 F D L A E D A K L Q L S K D T Y D D D L 300
901 gacaacctcctggcccagatcggcgaccagtacgccgacctgttcctcgcagccaagaac 960
301 D N L L A Q I G D Q Y A D L F L A A K N 320
961 ctgagcgacgccatcctcctgagcgacatcctcagagtgaacaccgagatcaccaaggct 1020
321 L S D A I L L S D I L R V N T E I T K A 340
1021 cccctgagcgccagcatgatcaagagatacgacgagcaccatcaggacctgaccctcctg 1080
341 P L S A S M I K R Y D E H H Q D L T L L 360
1081 aaggccctcgtgagacaacagctgcccgagaagtacaaggagatcttctttgaccagagc 1140
361 K A L V R Q Q L P E K Y K E I F F D Q S 380
1141 aagaacggctacgccggctacatcgacggaggcgccagtcaggaagagttctacaagttc 1200
381 K N G Y A G Y I D G G A S Q E E F Y K F 400
1201 atcaagcccatcctggagaagatggacggcaccgaagagctgctcgtgaagctgaacaga 1260
401 I K P I L E K M D G T E E L L V K L N R 420
1261 gaggacctgctcagaaagcagagaaccttcgacaacggcagcatcccccaccagatccac 1320
421 E D L L R K Q R T F D N G S I P H Q I H 440
1321 ctgggcgagctgcacgccatcctgagacggcaggaggacttctaccccttcctgaaggac 1380
441 L G E L H A I L R R Q E D F Y P F L K D 460
1381 aacagagagaagattgaaaagatcctgaccttcagaatcccctactatgtgggccccctg 1440
461 N R E K I E K I L T F R I P Y Y V G P L 480
1441 gccagaggcaacagcagattcgcctggatgaccaggaagagcgaagagacaatcacacct 1500

481 A R G N S R F A W M T R K S E E T I T P 500
1501 tggaacttcgaagaggtggtcgacaaaggcgccagcgcccagagcttcatcgagagaatg 1560
501 W N F E E V V D K G A S A Q S F I E R M 520
1561 accaacttcgacaagaacctgcccaacgagaaggtgctgcccaagcacagcctcctgtac 1620
521 T N F D K N L P N E K V L P K H S L L Y 540
1621 gagtacttcaccgtgtacaacgagctgaccaaggtgaagtacgtgaccgagggcatgaga 1680
541 E Y F T V Y N E L T K V K Y V T E G M R 560
1681 aagcctgcctttctgagtggcgagcagaagaaagccatcgtggacctgctcttcaagacc 1740
561 K P A F L S G E Q K K A I V D L L F K T 580
1741 aacagaaaagtgaccgtgaagcagctgaaggaggactacttcaagaaaatcgagtgcttc 1800
581 N R K V T V K Q L K E D Y F K K I E C F 600
1801 gacagcgtggagatcagcggcgtggaggacagattcaacgccagcctgggcacctaccac 1860
601 D S V E I S G V E D R F N A S L G T Y H 620
1861 gacctgctcaagattatcaaagacaaggacttcctggacaacgaagagaacgaggacatc 1920
621 D L L K I I K D K D F L D N E E N E D I 640
1921 ctggaggacatcgtgctgacactgaccctcttcgaggacagagagatgatcgaagagaga 1980
641 L E D I V L T L T L F E D R E M I E E R 660
1981 ctgaagacctacgcccacctgttcgatgacaaggtgatgaagcagctgaagagacggaga 2040
661 L K T Y A H L F D D K V M K Q L K R R R 680
2041 tacaccggctggggcagactgagcagaaagctgatcaacggcatcagagacaagcagagc 2100
681 Y T G W G R L S R K L I N G I R D K Q S 700
2101 ggcaagaccatcctggacttcctgaagagcgacggcttcgccaacagaaacttcatgcag 2160
701 G K T I L D F L K S D G F A N R N F M Q 720
2161 ctgatccacgatgacagcctgaccttcaaggaggacatccagaaagcccaagtgagcggg 2220
721 L I H D D S L T F K E D I Q K A Q V S G 740
2221 cagggcgacagcctgcacgagcatatcgccaacctggctggcagccccgccatcaagaaa 2280
741 Q G D S L H E H I A N L A G S P A I K K 760
2281 ggcatcctgcagaccgtgaaggtcgtggacgagctggtcaaggtgatgggcagacacaag 2340
761 G I L Q T V K V V D E L V K V M G R H K 780
2341 cccgagaacatcgtgattgagatggccagagagaaccagacaacccagaagggccagaag 2400
781 P E N I V I E M A R E N Q T T Q K G Q K 800
2401 aacagcagagagagaatgaagagaatcgaagagggcatcaaggagctgggcagccagatc 2460
801 N S R E R M K R I E E G I K E L G S Q I 820
2461 ctgaaggagcaccccgtggagaacacccagctgcagaacgagaagctgtacctgtattac 2520
821 L K E H P V E N T Q L Q N E K L Y L Y Y 840
2521 ctgcagaacggcagagacatgtacgtggaccaggagctggacatcaacagactgagcgat 2580
841 L Q N G R D M Y V D Q E L D I N R L S D 860
2581 tacgacgtggatcacatcgtcccccagagcttcctgaaggatgacagcatcgataacaag 2640
861 Y D V D H I V P Q S F L K D D S I D N K 880
2641 gtgctgaccagaagcgacaagaacagaggcaagagcgacaacgtgcccagcgaagaggtc 2700
881 V L T R S D K N R G K S D N V P S E E V 900
2701 gtgaagaaaatgaagaactactggagacagctcctgaacgccaagctgatcacccagaga 2760
901 V K K M K N Y W R Q L L N A K L I T Q R 920
2761 aagttcgacaacctgaccaaggccgagagaggcgggctgagcgagctcgacaaagccggc 2820
921 K F D N L T K A E R G G L S E L D K A G 940
2821 ttcatcaagagacagctggtggaaaccagacagatcaccaagcacgtggcccagatcctg 2880
941 F I K R Q L V E T R Q I T K H V A Q I L 960
2881 gacagcagaatgaacaccaagtacgacgagaacgataagctgatcagagaggtgaaggtc 2940
961 D S R M N T K Y D E N D K L I R E V K V 980
2941 atcaccctgaagagcaaactggtgagcgacttcagaaaggacttccagttctacaaggtg 3000
981 I T L K S K L V S D F R K D F Q F Y K V 1000
3001 agagagatcaataactaccatcacgctcatgacgcctacctgaacgccgtcgtgggcacc 3060
1001 R E I N N Y H H A H D A Y L N A V V G T 1020
3061 gccctgatcaagaaataccccaagctggagagcgagttcgtgtacggcgactacaaggtg 3120

1021 A L I K K Y P K L E S E F V Y G D Y K V 1040
3121 tacgacgtgagaaagatgatcgccaagagcgagcaggagatcggcaaggccaccgccaag 3180
1041 Y D V R K M I A K S E Q E I G K A T A K 1060
3181 tacttcttttacagcaacatcatgaacttctttaagaccgagatcaccctggccaacggc 3240
1061 Y F F Y S N I M N F F K T E I T L A N G 1080
3241 gagatcagaaagaggcccctgatcgaaaccaacggcgaaaccggcgagatcgtgtgggac 3300
1081 E I R K R P L I E T N G E T G E I V W D 1100
3301 aagggcagagacttcgccaccgtgagaaaggtgctgagcatgccccaggtgaacatcgtg 3360
1101 K G R D F A T V R K V L S M P Q V N I V 1120
3361 aagaaaaccgaggtgcagaccggaggcttcagcaaggagagcatcctgcccaagagaaac 3420
1121 K K T E V Q T G G F S K E S I L P K R N 1140
3421 agcgacaagctgatcgccagaaagaaagactgggaccccaagaaatacggaggcttcgac 3480
1141 S D K L I A R K K D W D P K K Y G G F D 1160
3481 agccccaccgtggcctacagcgtgctggtcgtggccaaggtggagaagggcaagagcaag 3540
1161 S P T V A Y S V L V V A K V E K G K S K 1180
3541 aaactgaagagcgtgaaggagctcctgggcatcaccatcatggagagatccagcttcgag 3600
1181 K L K S V K E L L G I T I M E R S S F E 1200
3601 aagaaccccatcgacttcctggaggccaagggctacaaggaggtgaagaaagacctgatt 3660
1201 K N P I D F L E A K G Y K E V K K D L I 1220
3661 atcaagctgcccaagtacagcctgttcgagctggagaacggcagaaagagaatgctggcc 3720
1221 I K L P K Y S L F E L E N G R K R M L A 1240
3721 agcgccggcgagctgcagaagggcaacgagctggccctgcccagcaagtacgtgaacttc 3780
1241 S A G E L Q K G N E L A L P S K Y V N F 1260
3781 ctgtacctggccagccactacgagaagctgaagggcagccccgaggacaacgagcagaag 3840
1261 L Y L A S H Y E K L K G S P E D N E Q K 1280
3841 cagctgttcgtggagcagcacaagcactacctggacgaaatcattgagcagatcagcgag 3900
1281 Q L F V E Q H K H Y L D E I I E Q I S E 1300
3901 ttcagtaagagagtgatcctggctgacgccaacctggacaaggtgctgagcgcctacaac 3960
1301 F S K R V I L A D A N L D K V L S A Y N 1320
3961 aagcacagagacaagcccatcagagagcaggccgagaacatcattcacctgttcaccctg 4020
1321 K H R D K P I R E Q A E N I I H L F T L 1340
4021 accaacctgggcgcacccgcagccttcaagtacttcgacaccacaatcgacagaaagaga 4080
1341 T N L G A P A A F K Y F D T T I D R K R 1360
4081 tacaccagcaccaaggaggtgctggacgccaccctgatccaccagagcatcaccggcctg 4140
1361 Y T S T K E V L D A T L I H Q S I T G L 1380
4141 tacgaaaccagaatcgacctgtcacagcttgggggtgacggatcccctcctcccaagagg 4200
1381 Y E T R I D L S Q L G G D G S **P P P K R** 1400
4201 cccaggctggactaa 4215
1401 **P R L D** * 1404

**Supplementary Fig. 1: Representative indel spectrum and quality control diagrams for each *Cas9* mRNA used in the cell culture assay.** Indel spectrum and quality control diagram obtained from TIDE and ICE analyses for *JDS246-Cas9*, *GeneArt® CRISPR nuclease* and *heiCas9*. Note decreased number of wildtype alleles (grey dashed line in TIDE analysis) in *heiCas9* transfected cells and increased abundance of 26 nt deletion (black arrowhead in ICE analysis).

Fig. S1

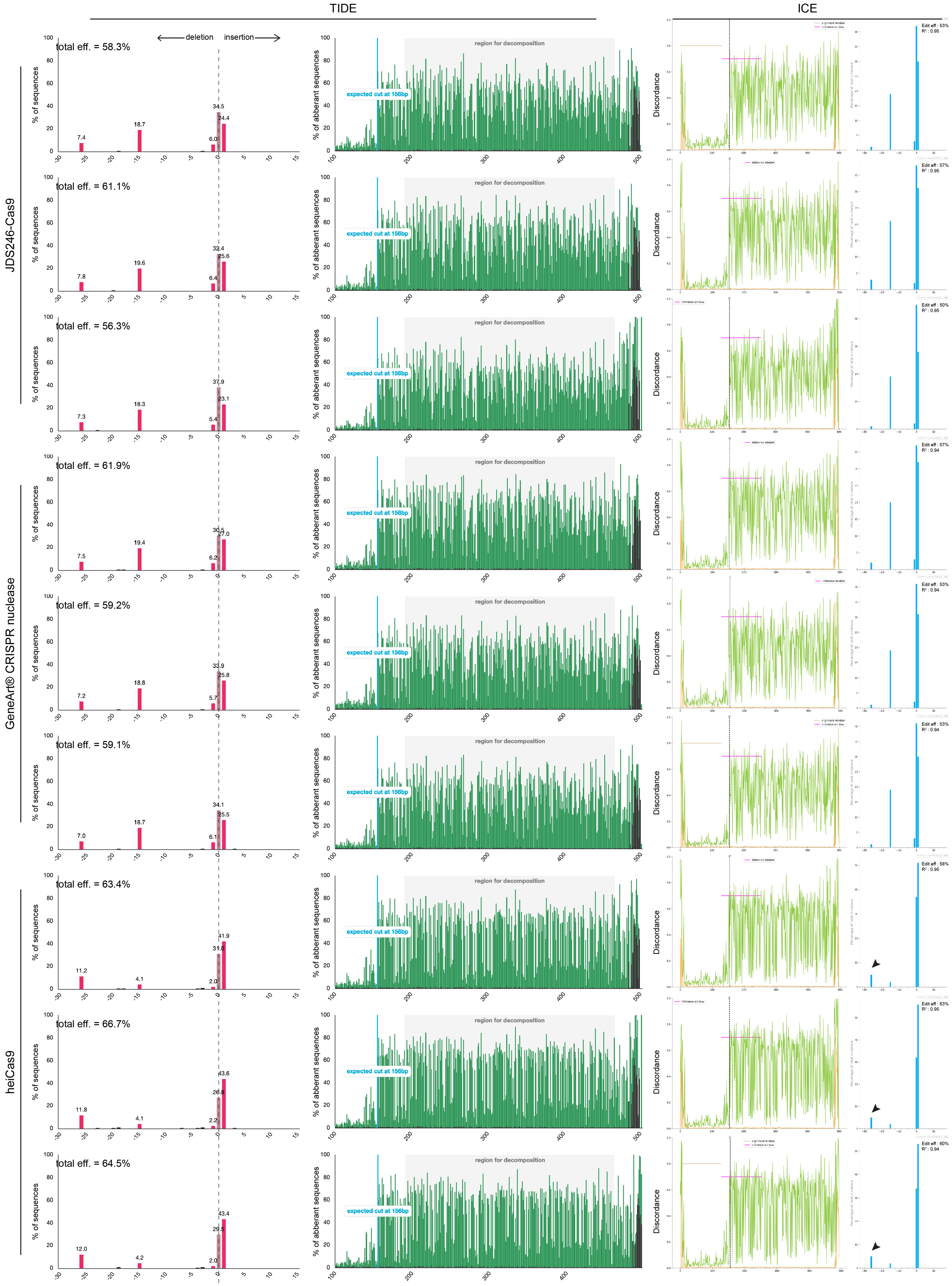
TIDE
ICE
JDS246-Cas9
GeneArt® CRISPR nuclease
heiCas9
deletion
insertion
% of sequences
% of aberrant sequences
Discordance
region for decomposition
expected cut at 156bp
total eff. = 58.3%
total eff. = 61.1%
total eff. = 56.3%
total eff. = 61.9%
total eff. = 59.2%
total eff. = 59.1%
total eff. = 63.4%
total eff. = 66.7%
total eff. = 64.5%
Edit eff : 53%
R² : 0.95
Edit eff : 57%
R² : 0.95
Edit eff : 50%
R² : 0.95
Edit eff : 57%
R² : 0.94
Edit eff : 53%
R² : 0.94
Edit eff : 53%
R² : 0.94
Edit eff : 58%
R² : 0.95
Edit eff : 63%
R² : 0.95
Edit eff : 60%
R² : 0.94

**Supplementary Fig. 2: Increased cytosine to thymine transition in medaka embryo pools injected with *heiBE4-Gam*.**

(a) Schematic representation of base editing window in *OlOca2 T1* target site.

(b-c) Sanger sequencing quantifications (EditR[28]) of pools of five randomly picked embryos injected with sgRNA *OlOca2 T1* and either *BE4-Gam* (b) or *heiBE4-Gam* (c). Note: in *heiBE4-Gam* injections, for each cytosine, the C-to-T transition rate was higher than 60%, a level never observed in *BE4-Gam* injected embryos. C997T is highlighted with white frame.

Fig. S2

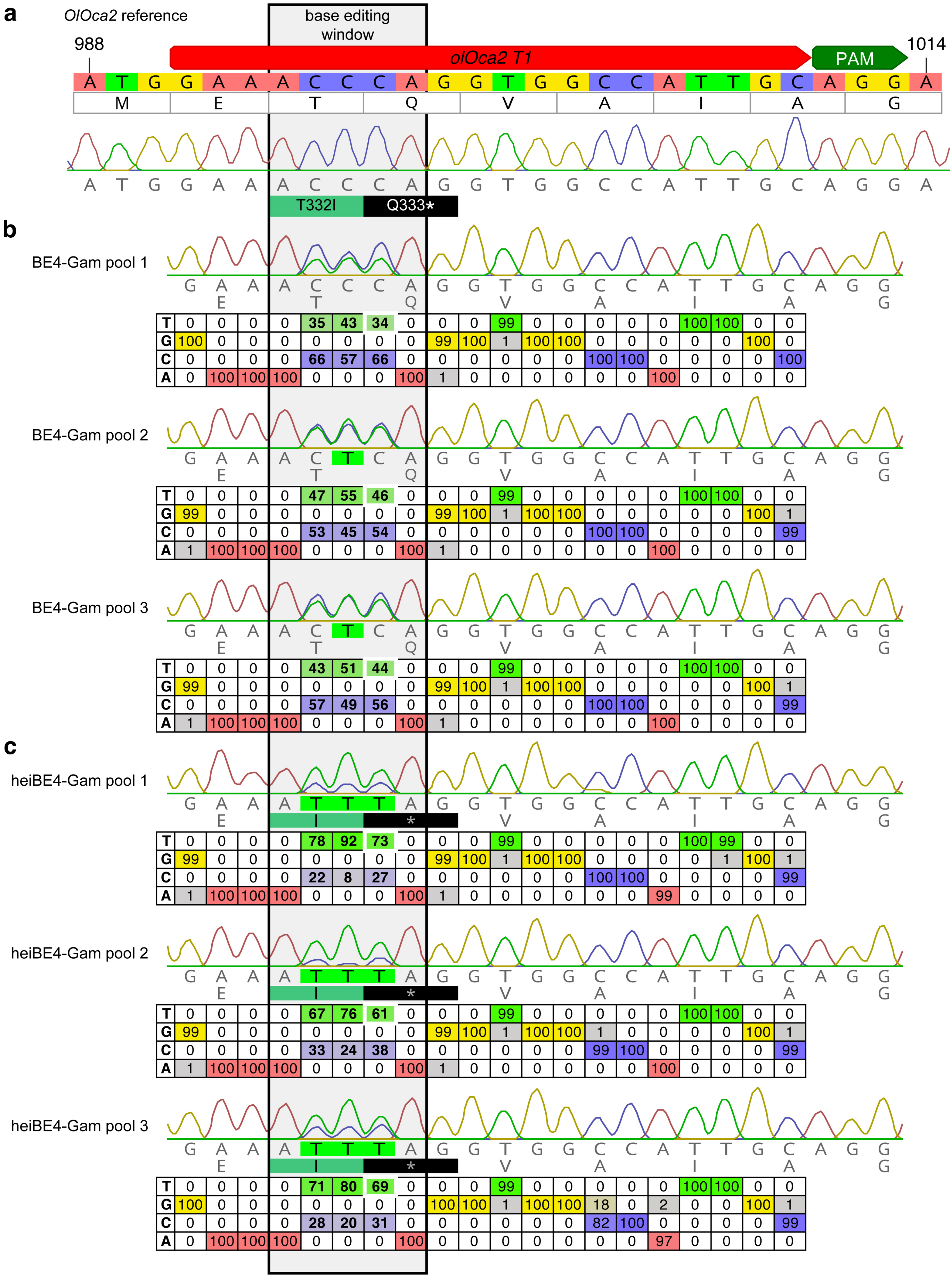


**b** BE4-Gam pool 1

| | | | | | | | | | | | | | | | | | | | | |
|---|---|---|---|---|---|---|---|---|---|---|---|---|---|---|---|---|---|---|---|---|
| T | 0 | 0 | 0 | 0 | 35 | 43 | 34 | 0 | 0 | 0 | 99 | 0 | 0 | 0 | 0 | 0 | 100 | 100 | 0 | 0 |
| G | 100 | 0 | 0 | 0 | 0 | 0 | 0 | 0 | 99 | 100 | 1 | 100 | 100 | 0 | 0 | 0 | 0 | 0 | 100 | 0 |
| C | 0 | 0 | 0 | 0 | 66 | 57 | 66 | 0 | 0 | 0 | 0 | 0 | 0 | 100 | 100 | 0 | 0 | 0 | 0 | 100 |
| A | 0 | 100 | 100 | 100 | 0 | 0 | 0 | 100 | 1 | 0 | 0 | 0 | 0 | 0 | 0 | 100 | 0 | 0 | 0 | 0 |

BE4-Gam pool 2

| | | | | | | | | | | | | | | | | | | | | |
|---|---|---|---|---|---|---|---|---|---|---|---|---|---|---|---|---|---|---|---|---|
| T | 0 | 0 | 0 | 0 | 47 | 55 | 46 | 0 | 0 | 0 | 99 | 0 | 0 | 0 | 0 | 0 | 100 | 100 | 0 | 0 |
| G | 99 | 0 | 0 | 0 | 0 | 0 | 0 | 0 | 99 | 100 | 1 | 100 | 100 | 0 | 0 | 0 | 0 | 0 | 100 | 1 |
| C | 0 | 0 | 0 | 0 | 53 | 45 | 54 | 0 | 0 | 0 | 0 | 0 | 0 | 100 | 100 | 0 | 0 | 0 | 0 | 99 |
| A | 1 | 100 | 100 | 100 | 0 | 0 | 0 | 100 | 1 | 0 | 0 | 0 | 0 | 0 | 0 | 100 | 0 | 0 | 0 | 0 |

BE4-Gam pool 3

| | | | | | | | | | | | | | | | | | | | | |
|---|---|---|---|---|---|---|---|---|---|---|---|---|---|---|---|---|---|---|---|---|
| T | 0 | 0 | 0 | 0 | 43 | 51 | 44 | 0 | 0 | 0 | 99 | 0 | 0 | 0 | 0 | 0 | 100 | 100 | 0 | 0 |
| G | 99 | 0 | 0 | 0 | 0 | 0 | 0 | 0 | 99 | 100 | 1 | 100 | 100 | 0 | 0 | 0 | 0 | 0 | 100 | 1 |
| C | 0 | 0 | 0 | 0 | 57 | 49 | 56 | 0 | 0 | 0 | 0 | 0 | 0 | 100 | 100 | 0 | 0 | 0 | 0 | 99 |
| A | 1 | 100 | 100 | 100 | 0 | 0 | 0 | 100 | 1 | 0 | 0 | 0 | 0 | 0 | 0 | 100 | 0 | 0 | 0 | 0 |

**c** heiBE4-Gam pool 1

| | | | | | | | | | | | | | | | | | | | | |
|---|---|---|---|---|---|---|---|---|---|---|---|---|---|---|---|---|---|---|---|---|
| T | 0 | 0 | 0 | 0 | 78 | 92 | 73 | 0 | 0 | 0 | 99 | 0 | 0 | 0 | 0 | 0 | 100 | 99 | 0 | 0 |
| G | 99 | 0 | 0 | 0 | 0 | 0 | 0 | 0 | 99 | 100 | 1 | 100 | 100 | 0 | 0 | 0 | 0 | 1 | 100 | 1 |
| C | 0 | 0 | 0 | 0 | 22 | 8 | 27 | 0 | 0 | 0 | 0 | 0 | 0 | 100 | 100 | 0 | 0 | 0 | 0 | 99 |
| A | 1 | 100 | 100 | 100 | 0 | 0 | 0 | 100 | 1 | 0 | 0 | 0 | 0 | 0 | 0 | 99 | 0 | 0 | 0 | 0 |

heiBE4-Gam pool 2

| | | | | | | | | | | | | | | | | | | | | |
|---|---|---|---|---|---|---|---|---|---|---|---|---|---|---|---|---|---|---|---|---|
| T | 0 | 0 | 0 | 0 | 67 | 76 | 61 | 0 | 0 | 0 | 99 | 0 | 0 | 0 | 0 | 0 | 100 | 100 | 0 | 0 |
| G | 99 | 0 | 0 | 0 | 0 | 0 | 0 | 0 | 99 | 100 | 1 | 100 | 100 | 1 | 0 | 0 | 0 | 0 | 100 | 1 |
| C | 0 | 0 | 0 | 0 | 33 | 24 | 38 | 0 | 0 | 0 | 0 | 0 | 0 | 99 | 100 | 0 | 0 | 0 | 0 | 99 |
| A | 1 | 100 | 100 | 100 | 0 | 0 | 0 | 100 | 1 | 0 | 0 | 0 | 0 | 0 | 0 | 100 | 0 | 0 | 0 | 0 |

heiBE4-Gam pool 3

| | | | | | | | | | | | | | | | | | | | | |
|---|---|---|---|---|---|---|---|---|---|---|---|---|---|---|---|---|---|---|---|---|
| T | 0 | 0 | 0 | 0 | 71 | 80 | 69 | 0 | 0 | 0 | 99 | 0 | 0 | 0 | 0 | 0 | 100 | 100 | 0 | 0 |
| G | 100 | 0 | 0 | 0 | 0 | 0 | 0 | 0 | 100 | 100 | 1 | 100 | 100 | 18 | 0 | 2 | 0 | 0 | 100 | 1 |
| C | 0 | 0 | 0 | 0 | 28 | 20 | 31 | 0 | 0 | 0 | 0 | 0 | 0 | 82 | 100 | 0 | 0 | 0 | 0 | 99 |
| A | 0 | 100 | 100 | 100 | 0 | 0 | 0 | 100 | 0 | 0 | 0 | 0 | 0 | 0 | 0 | 97 | 0 | 0 | 0 | 0 |
